# Random survival forests identify myocardial gene signatures associated with survival in heart failure with preserved ejection fraction

**DOI:** 10.1101/2025.06.08.658488

**Authors:** Suraj Kannan, Hildur Knutsdottir, Joban Vaishnav, Joel S. Bader, David A. Kass, Kavita Sharma, Virginia S. Hahn

## Abstract

Heart failure with preserved ejection fraction (HFpEF) continues to be poorly understood at the molecular level. While previous studies have identified gene expression signatures unique to HFpEF, genes associated with clinical decompensation have not been determined. Here, we performed exploratory analysis of myocardial RNA-seq data to identify genes associated with event-free survival in HFpEF. We analyzed previously published RNA sequencing data of right ventricular septal endomyocardial biopsies from HFpEF patients (n=41) with paired clinical, echocardiographic, and outcome data (including mortality and heart failure hospitalizations). We constructed random survival forests with forward stepwise regularization (the “variable hunting” method) to determine genes associated with time to first event using a combined end-point of heart failure hospitalization or all-cause death. Selection of candidate forest variables was tested with both random and weighted sampling methods. We identified 33 genes that are predictive of survival in HFpEF. This set includes genes previously implicated in heart failure, including ADAMTSL2, ADRB1, BMP6, and METRNL. Survival forests constructed using these genes outperform those constructed from clinical and hemodynamic parameters alone (out-of-bag C-index 0.894 vs 0.545). Moreover, in survival forests constructed from both gene data and hemodynamic measurements, individual survival genes consistently showed higher variable importance than clinical/hemodynamic parameters. Our study shows that random survival forests identify myocardial gene signatures that may better model HFpEF prognosis than clinical measurements. Further studies are warranted to validate findings in independent cohorts.

---

**T**hough several recent clinical trials have improved our therapeutic armamentarium for heart failure with preserved ejection fraction (HFpEF), improved understanding of the myocardial pathobiology is needed. Furthermore, identification of molecular phenotypes associated with worse survival could help identify novel or repurposed therapies. We previously reported a cohort of prospectively-defined HFpEF patients (n = 41) with paired endomyocardial biopsy RNA sequencing (RNA-seq), echocardiographic, hemodynamic, and outcome data (1). With this, we identified gene expression signatures unique to HFpEF. However, genes associated with clinical decompensation have not been determined.

Survival data is most commonly analyzed with methods based on Cox regression. These approaches typically rely on proportional hazards assumptions, and may struggle with modeling non-linear effects. Random survival forests (RSF), an extension of random forests to right-censored data, offer an alternative (2). Through bootstrap-aggregating, RSF can model complex survival functions with non-linear effects and interactions. In combination with stepwise regularization, RSF can be applied to ultra-high dimensional settings such as transcriptomic data (3). Here, we used RSF with the previously published HFpEF RNA-seq data to identify genes predictive of survival in HFpEF.

Our full methods, including reproducible code and rationale for parameters, can be found online (https://github.com/skannan4/hfpef_survival). We first constructed RSF with forward stepwise regularization (the “variable hunting” method (3)) to identify genes associated with time-to-first-event using a combined end-point of all-cause death or heart failure hospitalization. We selected genes (n = 4000) with the highest variance across HFpEF samples, using optimized forest parameter settings (ntrees = 1000, nodesize = 1; 500 total variable-hunting iterations). We tested four methods for weighting genes (p-values from initial differential expression testing, hazard ratios from simple Cox regression, variable importance in a test random forest, or uniform weighting). We then selected only the 33 genes that were identified by a majority of the methods to be predictive of survival **(Figure 1A)**.

**Fig. 1.**
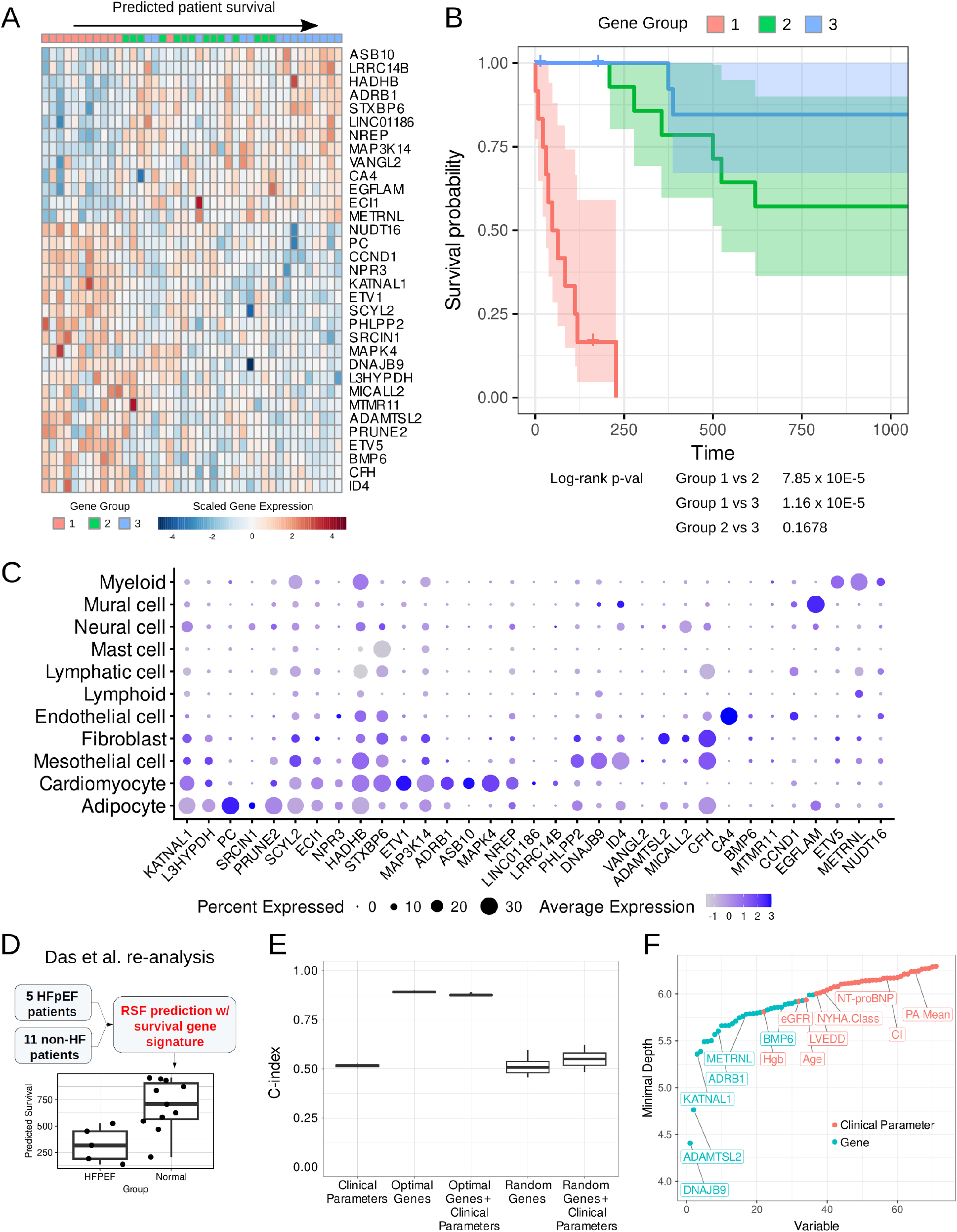
Random survival forests identify 33 genes predictive of HFpEF survival. **A**. Gene expression data for 33 survival genes (rows) plotted for 41 patients with HFpEF (columns) ordered by patient survival. Heatmap is annotated with the HFpEF subgroups categorized based on the expression of the 33 survival genes. **B**. Kaplan-Meier curves for three groups of patients clustered based on their expression of the survival genes. Log-rank p-values adjusted for sex, age, and eGFR. **C**. Expression of the 33 survival genes in snRNA-seq data taken from adult myocardium. **D**. Cross-validation of survival genes using an additional dataset of HFpEF samples. **E**. Performance of various combinations of genes and clinical parameters of HFpEF survival based on concordance index. **F**. Predictive importance of survival genes and clinical/hemodynamic parameters. Variable importance was assessed using the minimal depth method developed in (3).

To further validate, we categorized the HFpEF patients into three gene groups based on expression of the survival gene signature **(Figure 1B)**. Gene Group 1 had the worst survival compared to Gene Groups 2 and 3 (log rank p-values 7.85 *×* 10^−5^ for Gene Group 1 vs 2, 1.16 *×* 10^−5^ Group 1 vs 3, adjusted for age, sex, and renal function). Survival was not significantly different between Gene Groups 2 and 3 (adjusted log-rank p-value 0.17). The identified Gene Groups showed significant overlap with the previously published HFpEF subgroups (1). Seven of the 11 patients in the original HFpEF subgroup 1 (with the lowest event-free survival) were in Gene Group 1, while only 1 of 10 in the original HFpEF Subgroup 2 was in Gene Group 1 (Fisher’s exact p-value 0.024).

Our identified set included genes previously implicated in heart failure, including *ADAMTSL2* (negative regulator of TGF*β* signaling), *ADRB1* (*β*1 adrenergic receptor), *BMP6, KATNAL1* (involved in microtubule processing), and *METRNL* (adipokine protective against cardiac fibrosis). We investigated expression of these genes in an adult cardiac single-nuclei RNA-seq atlas **(Figure 1C)**. Interestingly, in addition to cardiomyocytes, many survival genes showed high expression in adipocytes, which is intriguing given the hypothesized role of epicardial fat in HFpEF pathobiology.

Cross-validation of our survival signature was limited by availability of other high-quality HFpEF RNA-seq datasets, particularly with paired survival data. As a proof-of-concept, we re-analyzed data from Das et al. (4) In this study, myocardial biopsy tissue RNA-seq was performed for 16 patients undergoing elective CABG, 5 of whom were prospectively classified as HFpEF-proxy. We found that RSF using our survival genes predicted lower survival for patients in the HFpEF group than the control group **(Figure 1D)**. This should be interpreted cautiously, as there was no survival data for the patients in the HFpEF group. Moreover, the control group did not represent patients with true lack of cardiac pathology. Despite these limitations, this analysis served as preliminary cross-validation of our identified survival genes.

We next sought to compare our gene signature with established clinical parameters associated with survival in HFpEF (5, 6) (including left ventricular end diastolic diameter, pulmonary artery mean pressure, cardiac index, and NT-proBNP; see Github for full list). Surprisingly, RSF using the survival genes outperformed RSF constructed from clinical and hemodynamic parameters alone (out-of-bag C-index 0.894 vs 0.545, **Figure 1E**). The addition of clinical parameters to the survival genes did not further improve prediction. In RSF constructed with clinical parameters and survival genes, survival genes had lower minimal depth, an indicator of variable predictive importance (3) **(Figure 1F)**. This may be partly due to the limited variability in the clinical parameters tested among HFpEF patients, or simply the improved prognostic yield of transcriptomics over clinical parameters. Interestingly, *NPPB* did not emerge in our final gene signature, and our identified genes outperformed serum NT-proBNP in terms of predictive importance despite its known prognostic role in HFpEF (7).

In summary, we used RSF to identify a gene signature associated with survival in HFpEF, which outperformed clinical parameters. Our results suggest that myocardial tissue-based -omics may better identify molecular phenotypes with worse survival than clinical phenotypes alone. Moreover, our gene signature may enable the identification of new biomarkers for HFpEF. Our dataset was limited both by sample size as well as having measurements only obtained at one static timepoint. It is possible that changes in parameters over time may be more illustrative than their starting value. There may thus be benefit to obtaining multiple measurements, both gene expression and clinical, from patients over time. Nevertheless, our study represents a first effort to identify gene expression signatures predictive of decompensation in HFpEF. Future studies are warranted to validate our findings in independent cohorts.

